# Neuroprotective role of lactate in a human *in vitro* model of the ischemic penumbra

**DOI:** 10.1101/2023.07.28.550936

**Authors:** Marta Cerina, Marloes Levers, Jason M. Keller, Monica Frega

## Abstract

In patients suffering from cerebral ischemic stroke, there is an urgent need for treatments to protect brain cells. Recently, treatment strategies that induce neuronal activity have been shown to be neuroprotective. However, the biological mechanisms underlying the benefit from neuronal activation are unknown. We hypothesized that neuronal activation might trigger the astrocyte-to-neuron lactate shuttle, whereby lactate is released from astrocytes to support the energy requirements of hypoxic neurons, and this leads to the observed neuroprotection. We tested this by establishing a human cell based *in vitro* model of the ischemic penumbra. We found that lactate transporters are involved in the neuroprotective effect mediated by neuronal activation, that lactate exogenously administered before hypoxia correlated with neuroprotection, and that stimulation of astrocyte with consequent endogenous production of lactate resulted in neuroprotection. We presented evidence that lactate contributes to neuroprotection during hypoxia providing a potential basis for therapeutic approaches in ischemic stroke.

## Introduction

Ischemic stroke is a pathological condition caused by the sudden interruption of blood flow to the brain. It is ranked as the second leading cause of death and a major cause of adult disability worldwide^1^. In the core of the infarct region, oxygen and glucose deprivation leads to an energy failure that disrupts ion gradients across the plasma membrane and impairs neuronal function, as well as induces cell swelling and death within minutes^2^. Surrounding the core is the ischemic penumbra, a region where some remaining blood flow from collateral vessels persists^3, 4^. This area is functionally silent but structurally intact and viable with the potential to recover without damage, but only if normal blood perfusion is restored in time^3, 4^. Otherwise, the tissue becomes progressively and irreversibly damaged^3, 4^. For this reason, the penumbra represents the most therapeutically relevant target in ischemic stroke research.

The only treatment proven to benefit patients is early removal of the arterial occlusion (i.e., acute recanalization)^5, 6^. However, due to strict selection criteria, only very few patients are eligible for this type of treatment and, even if successful, most experience persistent neurological deficits^5, 6^. Thus, alternative, or adjunct neuroprotective strategies counteracting the molecular and cellular events causing ischemic damage in the penumbra have been sought^7^. The aim is to slow down the pathophysiological cascade, promote neuronal survival, and support functional recovery and neurological improvement^8, 9^. Over the past two decades, animal studies have shown that strategies directed at suppressing neuronal activity to preserve energy for basic cellular functions protect brain tissue^10^. However, in clinical trials promising pre-clinical studies have not been translated into positive outcomes^11, 12^. Failure of translation to patients may be associated with the mechanism of action of treatments under study. In fact, contrary to the prevailing view on suppression, inhibiting neuronal activity during hypoxia has been associated with irreversible tissue damage both *in vitro* and in patients^13–16^. Alternatively, we have shown that neuronal activation has a beneficial effect in both rodent and human neuronal cell cultures exposed to hypoxic conditions^17, 18^. However, the biological mechanisms underlying the benefit from neuronal activation during hypoxia are unknown. The identification of such mechanisms would provide new insights for the development of neuroprotective strategies.

Although lactate has long been considered a by-product of anaerobic metabolism, recent evidence showed that it can be found within the extracellular space of human neuronal tissue under aerobic conditions^19^, suggesting that it could have a physiological role in the brain as a supplemental energy source or even as a signalling molecule^20^. In this context, Pellerin and Magistretti proposed an astrocyte-to-neuron lactate shuttle (ANLS) hypothesis where astrocytes that respond to neuronal activity (e.g., via glutamate signalling) release lactate into the extracellular space to energetically support neurons^21, 22^. Recent evidence also suggests that lactate might be a neuroprotective agent in certain pathological contexts. Studies in animal models of ischemic stroke showed that lactate, either endogenously produced during hypoxia or applied exogenously at reoxygenation, supports energy metabolism and functional recovery in neurons^23–26^, and that direct intracerebroventricular injection of lactate led to a decrease in infarct volume and an improvement in neurological outcome^26, 27^. Moreover, it has been demonstrated that lactate mediates neuroprotection through different mechanisms of action which are relevant in ischemic stroke (i.e., support to energy metabolism^23–25^, counteraction of glutamate excitotoxicity^27–29^, oxidative stress^28–30^ and cellular death^31, 32^, expression of plasticity-related and pro-survival genes ^31, 32^, and protection of astrocytes^33^). According to the ANLS hypothesis, lactate production and release by astrocytes is dependent on neuronal activity^21, 22^. Therefore, we hypothesised that neuronal activation might maintain or trigger the ANLS to protect hypoxic neurons.

Here, we established a human *in vitro* neuronal model of the ischemic penumbra to investigate whether lactate might be neuroprotective in this setting. We found that lactate transporters are critically involved in the neuroprotective effect mediated by neuronal activation. Furthermore, we show that lactate exogenously administered before hypoxia correlates with neuroprotection in our model. Finally, we observed that activation of astrocyte with consequent endogenous production of lactate is associated with neuroprotection.

## Results

### Neuronal activation protects cells during hypoxia

We differentiated hiPSCs into human neuronal networks (Fig. 1a) and investigated the effect of artificial activation during 24 hours of low oxygen exposure on neuronal activity and viability. By seven weeks *in vitro*, hiPSCs-derived neuronal networks grown on MEAs were functionally mature, showing typical neuronal morphology and stable electrophysiological activity composed of spikes and synchronous bursts^18, 34, 35^ (Fig. 1b). To ensure consistency between cell cultures, all hiPSCs-derived neuronal networks were used in hypoxia experiments at this stage of maturation^36^. When exposed to low oxygen, both the firing rate and network bursting activity of the networks progressively declined, leading to a complete inhibition of network bursts within 24 hours (Fig. 1c-e). In line with our previous results^18^, we found that when networks were artificially activated during hypoxia the MFR was maintained at higher levels compared to untreated neuronal networks (Fig. 1c-e). Additionally, the NBR was preserved at baseline levels for the first 18 hours of hypoxia (p > 0.999), and it was significantly higher at every timepoint compared to untreated neuronal networks (p < 0.0001 at each timepoint, Fig. 1d,e). Next, we evaluated the effect of neuronal activation on cell viability by determining the percentage of apoptotic and dead cells after 24 hours of hypoxia (Fig. 1f). Corroborating our MEA recordings, we found a significantly higher percentage of apoptotic cells in untreated neuronal networks compared to those where neuronal activation was applied during hypoxia (p < 0.05, Fig. 1g). However, the percentage of dead cells was not statistically different between the two experimental groups (p > 0.9999, Fig. 1g).

**Fig. 1.**
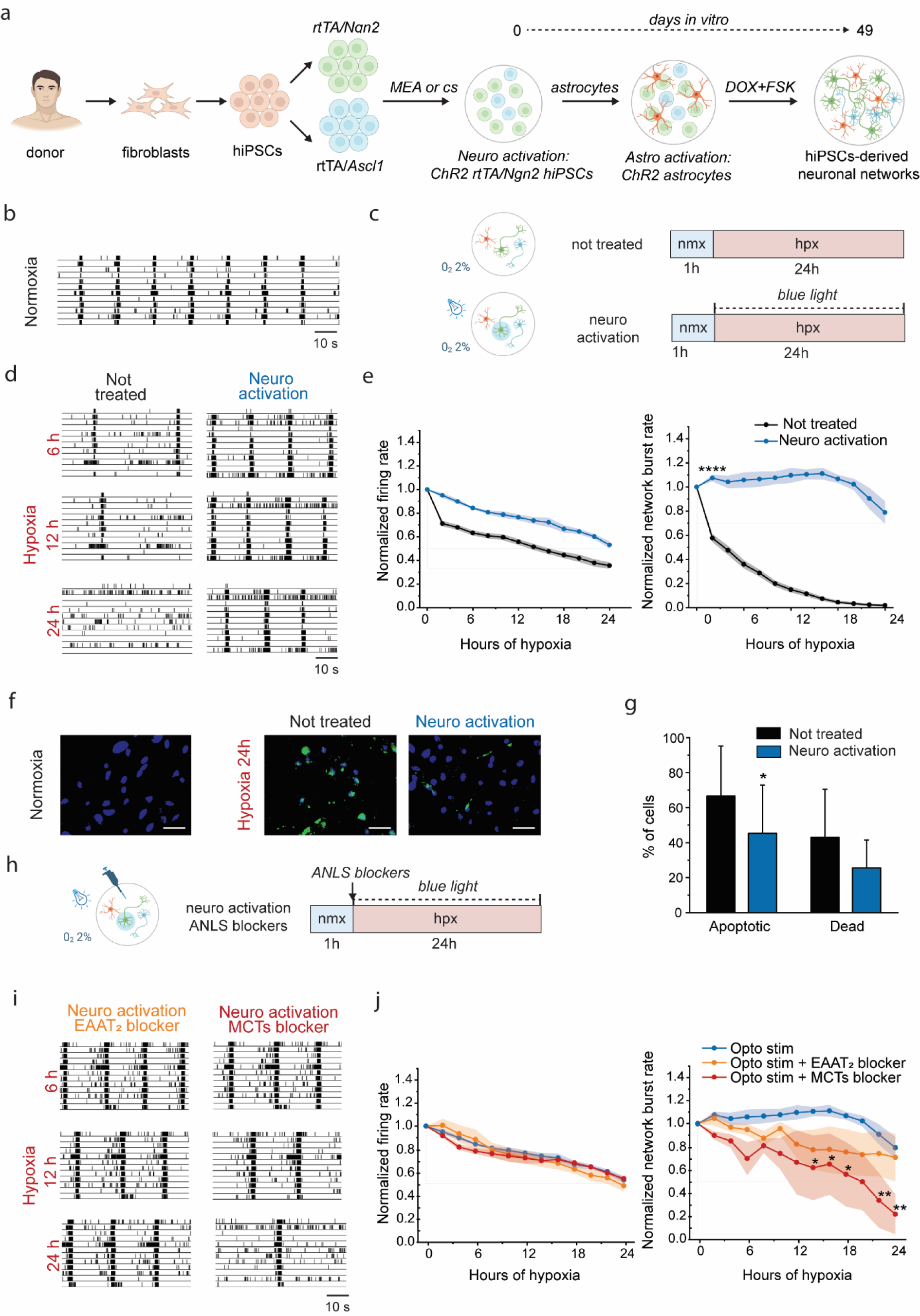
Lactate transport is involved in neuroprotection mediated by neuronal activation. **a** Schematic presentation of the differentiation protocol of hiPSCs reprogrammed from fibroblasts of healthy donors. **b** Representative raster plots showing 3 minutes of electrophysiological activity exhibited in normoxia by functionally mature healthy neuronal networks. **c** Schemes showing the experimental protocol, indicating the duration of normoxia (nmx, blue) and hypoxia (hpx, red), and experimental conditions during hypoxia (i.e., not treated and activated with optogenetic stimulation (neuro activation)). **d** Representative raster plots showing 60 seconds of electrophysiological activity exhibited at different timepoints of hypoxia (6, 12, 24 hours) by neuronal networks not treated and activated with optogenetic stimulation (neuro activation). **e** Graphs showing the effect of 24 hours of hypoxia on MFR and NBR in neuronal networks not treated and activated with optogenetic stimulation. The values are normalized to the data of normoxia (not treated n=20, neuro activation n=5). ****p<0.0001, two-way ANOVA test and post hoc Bonferroni correction was performed between conditions. Asterisks are not shown after the first timepoint the corresponding p-value was found. **f** Representative images of live/dead assay in normoxia and after 24 hours of hypoxia in neuronal networks not treated or activated with optogenetic stimulation. Neurons were stained for DAPI (blue, nuclei of all cells) and CellEvent™ Caspase-3/7 Green Detection Reagent (green, apoptotic cells). Scale bar: 20 μm. **g** Bar graphs showing the percentage of apoptotic cells and dead cells in hiPSCs-derived neurons not treated and activated with optogenetic stimulation (not treated n=2, neuro activation n=2). *p<0.05, Mann Whitney test was performed between conditions. **h** Scheme showing the experimental protocol, indicating the duration of normoxia (nmx, blue) and hypoxia (hpx, red), and experimental condition during hypoxia (i.e., activated with optogenetic stimulation with, alternatively, EAAT2 blocker or MCTs blocker (neuro activation + ANLS blockers)). **i** Representative raster plots showing 60 seconds of electrophysiological activity exhibited at different timepoints of hypoxia (6, 12, 24 hours) by neuronal networks activated with optogenetic stimulation with, alternatively, EAAT2 blocker or MCTs blocker. **j** Graphs showing the effect of ANLS blockers on MFR and NBR in neuronal networks activated with optogenetic stimulation. The values are normalized to the data of normoxia (neuro activation n=5, neuro activation + EAAT2 blocker n=5, neuro activation + MCT1/2 blocker n=3). *p<0.05, **p< 0.005, two-way ANOVA test and post hoc Bonferroni correction was performed between conditions.

### Lactate transporters are involved in neuroprotection mediated by neuronal activation during hypoxia

We turned our attention to the biological mechanisms underlying the neuroprotective effect that artificial activation has during hypoxia. To evaluate this, we pharmacologically blocked transporters that are predicted to be directly involved in the ANLS (Fig. 1h). In particular, we blocked excitatory amino acid transporter 2 (i.e., EAAT2 or GLT-1, responsible for glutamate uptake into astrocytes^22, 37^), and two monocarboxylate transporters (i.e., MCT1 and MCT2, the major lactate transporters present on neurons as well as glial cells^38^). The effects of the blockers were primarily observable on the NBR: inhibiting EAAT2 slightly impaired neuroprotection at the highest concentration tested (Fig. 1 i,j, Suppl. Fig. 1), whereas blocking MCT1/2 showed a strong inhibition of neuroprotection (p<0.05 after 14, 16 and 18 hours, p<0.005 after 22 and 24 hours, Fig. 1i,j).

### Exogenous lactate administrated before hypoxia is neuroprotective

Since blocking MCT1/2 impaired the neuroprotective effect mediated by neuronal activation, we postulated that lactate itself, whose production and release by astrocytes could be maintained or triggered by neuronal activation, may have a neuroprotective effect on its own in our model.

First, we tested this hypothesis by adding L-lactate (5 mM, 10 mM and 20 mM) to the culture medium of neuronal networks before exposing them to low oxygen (Fig. 2a). We found that L-lactate immediately blunted the electrophysiological activity of hiPSCs-derived neuronal networks before hypoxia onset, in a concentration-dependent manner (Suppl. Fig. 2a,b). Therefore, we decided to investigate the potential neuroprotective effect of L-lactate by evaluating the percentage of live, apoptotic and dead cells after 24 and 48 hours of hypoxia.

**Fig. 2.**
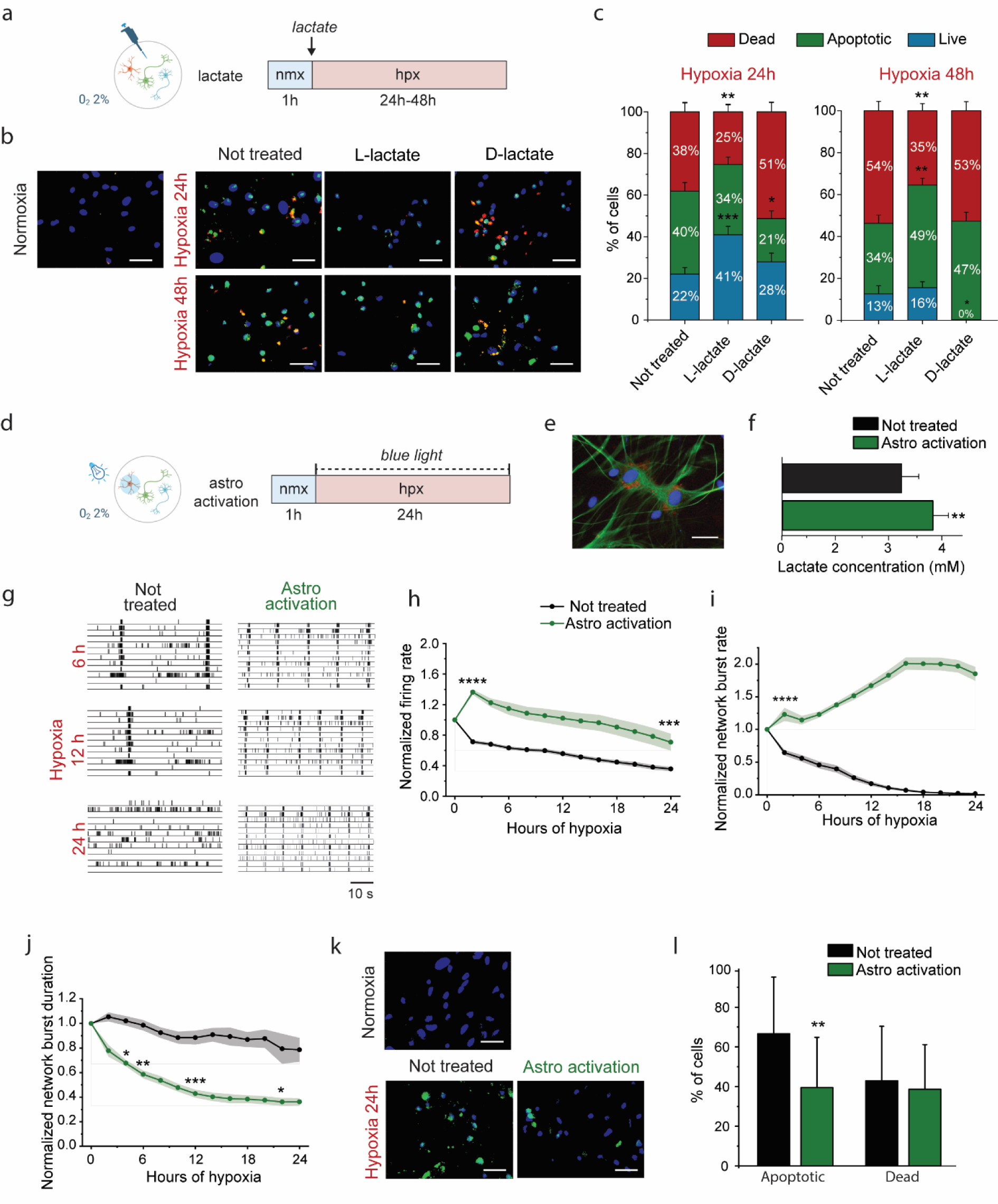
Lactate is neuroprotective in neuronal networks exposed to hypoxia. **a** Scheme showing the experimental protocol, indicating the duration of normoxia (nmx, blue) and hypoxia (hpx, red), and experimental condition during hypoxia (i.e., treated with lactate before hypoxia). **b** Representative images of live/dead assay in normoxia and after 24 and 48 hours of hypoxia in hiPSCs-derived neurons not treated and treated with 10 mM of L-lactate and 10 mM of D-lactate stained for DAPI (blue), CellEvent™ Caspase-3/7 Green Detection Reagent (green) and PI (red). Scale bar: 20 μm. **c** Stacked column graphs showing the percentage of live, apoptotic and dead cells (not treated n=7, L-lactate 10 mM n=5, D-lactate 10 mM n=2). *p<0.05, **p< 0.005, ***p<0.0005, unpaired t test or Mann Whitney test was performed between conditions. **d** Scheme showing the experimental protocol, indicating the duration of normoxia (nmx, blue) and hypoxia (hpx, red), and experimental condition during hypoxia (i.e., activation of astrocytes (astro activation)). **e** A representative image showing hiPSCs-derived neurons with rodent astrocytes stained for GFAP (green), cell nuclei stained with DAPI (blue), ChR2 (red). **f** Bar graph showing lactate concentration in the medium collected from astrocytic cultures not treated and activated with optogenetic stimulation. **g** Representative raster plots showing 60 seconds of electrophysiological activity exhibited at different timepoints of hypoxia (6, 12, 24 hours) by neuronal networks not treated and with activation of astrocytes. **h-j** Graphs showing the effect of 24 hours of hypoxia on MFR, NBR and NBD in neuronal networks not treated and with activation of astrocytes. The values are normalized to the data of normoxia (not treated n=20, astro activation n=9). *p<0.05, **p<0.005, ***p<0.0005, ****p<0.0001, two-way ANOVA test and post hoc Bonferroni correction was performed between conditions. Asterisks are not shown after the first timepoint the corresponding p-value was found. **k** Representative images of live/dead assay in normoxia and after 24 hours of hypoxia in neuronal networks not treated or with activation of astrocytes. Neurons were stained for DAPI (blue, nuclei of all cells) and CellEvent™ Caspase-3/7 Green Detection Reagent (green, apoptotic cells). Scale bar: 20 μm. **l** Bar graphs showing the percentage of apoptotic and dead cells in hiPSCs-derived neurons not treated and with activation of astrocytes (not treated n=2, astro activation n=2). **p<0.005, Mann Whitney test was performed between conditions.

After 24 hours of hypoxia, neuronal networks treated with 10 mM of L-lactate showed a higher percentage of live cells, plus a lower percentage of apoptotic and dead cells, as compared to neuronal networks exposed to hypoxia without lactate supplementation (Fig. 2b,c). Differences in the numbers of live and dead cells were statistically significant (p < 0.0005 and p < 0.005, respectively), whereas the quantity of apoptotic cells was not (p > 0.9999). After 48 hours of hypoxia, the percentage of live cells remaining in neuronal networks treated with L-lactate was very similar to untreated neuronal networks (p > 0.9999). However, the percentage of apoptotic cells was significantly higher (p < 0.005), and the percentage of dead cells was significantly lower (p < 0.005, Fig. 2b,c). These results suggest that adding 10 mM of L-lactate to the culture medium before hypoxia delayed apoptosis and consequent cell death. However, we found that lower or higher concentrations of L-lactate (i.e., 5 mM and 20 mM) had no measurable benefit on cell viability in our test system. While 5 mM of L-lactate did not have any deleterious or positive effects on the percentage of live cells, 20 mM of L-lactate appeared to be neurotoxic by increasing the number of apoptotic and dead cells (Suppl. Fig. 2c).

Next, to discriminate whether the potential neuroprotective effect of lactate depends on its modulatory effect of electrophysiological activity, on its use as a metabolic substrate, or on other mechanisms, we compared the effects of the addition of D-lactate (10 mM) and L-lactate. The D enantiomer of lactate, indeed, is poorly metabolized by neurons^39^, but it can be up taken by neurons via MCT1^38^ and is a partial agonist of the HCA1 receptor (i.e., responsible for the modulation of neuronal activity^40^). As was observed for L-lactate, D-lactate also immediately inhibited electrophysiological activity (Suppl. Fig. 2a,b). At 24 hours of hypoxia, we found a higher percentage of dead cells in neuronal networks treated with D-lactate compared to untreated neuronal networks (Fig. 2b,c). After 48 hours of hypoxia, the percentages of both apoptotic and dead cells in cultures treated with D-lactate were higher than in untreated neuronal networks, and the percentage of live cells was close to zero (Fig. 2b,c).

### Lactate is released by astrocyte activation and correlates with neuroprotection

We observed that the administration of lactate before hypoxia onset correlated with neuroprotection. Then, we decided to investigate whether lactate released by astrocytes during hypoxia had a similar effect.

To induce lactate release from astrocytes, we transduced the cells with ChR2 and stimulated them optogenetically with blue light^41^ (Fig. 2d). ChR2 integration was cell-specific and successful, as shown by ChR2 expression only in astrocytes (Fig. 2e). We found that the concentration of lactate was significantly higher in the medium of cultures underwent astrocyte activation (p<0.005, Fig. 2f), confirming that stimulation induced lactate release from astrocytes.

When exposed to 2% oxygen, we observed higher MFRs and NBRs in neuronal networks where astrocytes were activated optogenetically as compared to untreated cultures (p < 0.0001 during the first 22 hours of hypoxia and p<0.005 afterwards for MFR, p<0.0001 at each timepoint for NBR, Fig. 2g-i). Moreover, astrocyte activation shortened the NBD, as compared to not treated neuronal networks, at each timepoint during the hypoxic period (Fig. 3g,j).

Next, we investigated the effect of astrocyte activation on cell viability, by evaluating the percentage of apoptotic and dead cells after 24 hours of hypoxia. We found that the percentage of apoptotic cells in neuronal networks where astrocytes were activated was significantly lower compared to untreated networks (p < 0.05, Fig. 2k,l). However, the percentage of dead cells was not statistically different between the two test conditions (p > 0.9999, Fig. 2k,l).

## Discussion

In this study, we used hiPSCs-derived neuronal networks exposed to hypoxia as a human *in vitro* model of the ischemic penumbra to investigate whether the ANLS is involved in the neuroprotective effect mediated by neuronal activation. We found that the blockade of lactate transporters impairs the neuroprotective effect mediated by neuronal activation, clearly indicating that lactate transport is involved in neuroprotection. Furthermore, we observed that supplementing L-lactate, but not D-lactate, in the culture medium prior to the onset of hypoxia had a neuroprotective effect with respect to cell viability. This implies that lactate is neuroprotective in our model of the ischemic penumbra, and that this beneficial effect is mediated by the transport and metabolism of lactate in neurons. Lastly, we showed that optogenetic stimulation of astrocytes during hypoxia induced released of lactate and was correlated with neuroprotection, as demonstrated by the preservation of both spontaneous electrophysiological activity and cell viability.

We showed that the blockade of two transporters involved in the ANLS partially impairs the neuroprotective effect mediated by neuronal activation via optogenetic stimulation. We found that inhibiting EAAT2, which mediates glutamate uptake into astrocytes, a trigger for lactate production and release into the extracellular space^22, 37^, only mildly affected the optogenetic stimulation effect on synchronized activity in a concentration dependent manner. On the contrary, blocking MCT1/2, which mediate lactate transport at both neuronal and astrocytic level, showed a strong inhibition of neuroprotection. These results clearly showed that lactate transport via MCTs plays a critical role in neuroprotection mediated by neuronal activation in hypoxia, indicating that neuronal activation induced lactate release by astrocytes and/or lactate uptake by neurons.

Results obtained with the blockade of MCTs suggested that lactate in itself might be neuroprotective during hypoxia. Therefore, we decided to test this hypothesis by supplying neurons with lactate in two different ways: (i) by administering lactate before hypoxia onset, and (ii) by inducing lactate release through optogenetic stimulation of astrocytes. In both ways, we observed a neuroprotective effect in our model.

In line with previous results obtained in rodent neuronal cultures^40^, we found that the administration of lactate in a concentration range of 5-20 mM (which are relatively high as compared to physiological ones^19^) caused an immediate drop in the electrophysiological activity of hiPSCs-derived neuronal networks, in a concentration-dependent manner. To our knowledge, this is the first study demonstrating that lactate has a modulatory effect on activity of human neurons. Both L-lactate and D-lactate had the same effect on electrophysiological activity, since they modulate neuronal activity by binding the same receptor on the plasma membrane^40^. For this reason, we decided to assess the potential neuroprotective effect of L-lactate and D-lactate on cell viability. Our results showed that treatment with 10 mM of L-lactate induces a delay in apoptosis, and subsequent death, of neuronal cells exposed to hypoxia. Conversely, in neuronal networks treated with D-lactate we observed an increase in apoptosis and cell death as compared to not treated cultures. Since D-lactate can modulate neuronal activity but cannot be metabolized by neuronal cells, its failure as a neuroprotective agent allowed us to draw two conclusions. Firstly, the neuroprotective effect of L-lactate is not, or at least not only, dependent on the partial silencing of electrophysiological activity but depends on its use as a metabolic substrate. Secondly, D-lactate neurotoxic effect indicates that suppression of neuronal activity during hypoxia alone is not effective in protecting human neurons from ischemic damage, which is consistent with the above mentioned failure of neuroprotective strategies directed to suppression of neuronal activity during hypoxia^11, 12^. Moreover, because the MCTs carries both L-and D-lactate, the latter might act as a competitive inhibitor of L-lactate transport, and so reducing neuroprotection mediated by the uptake of L-lactate already present in the medium.

For a long time, lactate in the adult brain has not been considered more than a mere by-product of anaerobic metabolism. However, recent evidence suggested that lactate has a physiological role in the brain as a supplementary energy source and signaling molecule^20^, and a neuroprotective effect in different pathological conditions, including cerebral ischemic stroke. Studies on *in vitro* rodent models, showed that lactate, either endogenously produced during hypoxia or applied exogenously at the end of it, can be metabolically used by neurons, and it is preferential to glucose for functional recovery during the reoxygenation period^23–26^. In *in vivo* models, Schurr *et al*. found that the blockade of lactate transport exacerbates neuronal damage after ischemia, thus showing that endogenously produced lactate contributes to neuroprotection during global ischemia^42^. Berthet *et al*. showed that the intracerebroventricular injection of lactate after reperfusion led to a significant decrease in lesion size and an improvement in neurologic outcome in a rat model of cerebral ischemia^26, 27^.

According to literature, lactate contributes to neuroprotection through different mechanisms of action. Firstly, lactate represents a convenient metabolic substrate for energy-deprived neurons, since, as compared to glucose, its use reduces the strain on the depleted energy levels. Secondly, lactate counteracts glutamate excitotoxicity^43–45^, by triggering well-coordinated mechanisms leading to hyperpolarization of neurons, and thus to a decrease of neuronal excitability^44^, and increased expression of anti-apoptotic factors^46^. Furthermore, recent studies have investigated the neuroprotective effect of lactate against oxidative stress, showing that lactate triggers antioxidant defenses and pro-survival pathways, through the production of NADH, reduction of intracellular pH and a mild Reactive Oxygen Species (ROS) burst^28–30^. Thirdly, it has been demonstrated that physiological concentrations of lactate only, not glucose or pyruvate, promote the expression of key plasticity-related genes^31^ and that lactate promotes the upregulation of pro-survival genes and the downregulation of pro-death genes in an NMDARs-dependent manner, revealing a transcriptome profile favoring neuroprotection^32^.

Fourthly, lactate exerts its neuroprotective effect by targeting astrocytes. In particular, Benerjee *et al*. demonstrated that ischemic concentrations of lactate promote TWIK-related potassium channel (TREK1) expression, which contributes to maintain astrocytic properties (i.e., conductance, resting membrane potential, potassium buffering, glutamate uptake and pH), underlying ionic homeostasis and excitotoxicity prevention^33^.

Since the pathophysiological cascade of ischemic stroke is incredibly complex, a neuroprotective agent targeting more than one pathophysiological event could represent a successful approach to develop clinically relevant therapies for ischemic stroke. Actually, a few studies investigated treatment strategies with multiple therapeutic targets providing strong proof of principle of their effectiveness in neuroprotection in ischemic stroke^47–49^. In addition, we suggest that lactate represents a convenient neuroprotective agent in the perspective of translation to clinic since it can be administrated to patients in a simple way which is intravenous supplementation. In recent years, the administration of lactate enriched solutions has been studied in healthy subjects, and in acute heart failure and severe traumatic brain injured patients, showing beneficial effects, and no severe adverse event^50^.

Astrocyte activation during hypoxia, with the subsequent release of small concentrations of lactate in the extracellular space, was also associated to neuroprotection in our model. In particular, astrocyte stimulation resulted in the decrease of apoptotic and dead cells, as compared to not treated neuronal networks, and the preservation of electrophysiological activity throughout the 24 hours of hypoxia. The release of small concentrations of lactate over time had a very different effect as compared to the complete abolishment of activity observed when high concentrations of lactate were exogenously administered at once. In fact, with astrocyte activation the activity was finely modulated during hypoxia: we observed an increase in the frequency of network bursts, and a shortening of their duration. Both the balance AMPARs/NMDARs-mediated currents, and excitation/inhibition has been found to regulate the network burst activity in hiPSCs-derived neuronal networks^34, 35^. Since lactate potentiates NMDARs-mediated currents^31^ and has an inhibitory effect on neuronal activity^40^, a change in the frequency and duration of network bursts can be associated with the effect of lactate on neurons.

Even though we confirmed that astrocyte activation induces lactate release in the extracellular space, we cannot exclude that other mechanisms might be activated by optogenetic stimulation of astrocytes. In fact, astrocytes in the brain are involved in numerous critical functions in support of neuronal functionality and viability in both physiological and pathological contexts, including ischemic stroke^51–53^. During the acute phase of ischemic stroke, astrocytes undergo important morphological modifications and perform multiple functions both detrimental and beneficial for neuronal survival^54^. Within minutes after injury, activated astrocytes, also referred to as reactive astrogliosis, produce and release inflammatory mediators^55^ and ROS^56^, which may induce apoptosis and cell death. However, they also release neurotrophic factors in support of neurons^57^ and they contribute to neuroprotection against glutamate excitotoxicity^58^ and oxidative stress^59, 60^. During the late recovery phase after stroke, astrocytes participate in neurogenesis^61^, synaptogenesis^62, 63^, angiogenesis^64^, and thereby promote neurological recovery. For all these reasons, neuroprotective strategies that specifically target neurons alone may be insufficient to improve neurological outcome after ischemic stroke. In the past years, several strategies directed at reducing the detrimental effects and amplifying the beneficial effects of astrocytes on neuroprotection and neurorestoration have been investigated^65, 66^. Some of those approaches aimed, for instance, to protect astrocyte viability, enhance their ability to protect neurons from excitotoxicity and oxidative damage, and promote the release of neurotrophins and other factors for neuro- and angiogenesis^65, 66^. Developing treatment strategies that target multiple cell types, and in particular astrocytes, may represent a successful approach in ischemic stroke.

## Experimental procedures

### hiPSCs generation and neuronal differentiation

The hiPSCs used in this study were generated from fibroblast and kindly provided by Mossink et al^34^. Control line 1 is generated from a 30 year-old female and reprogrammed via episomal reprogramming (Coriell Institute for medical research, GM25256) and control line 2 was generated from a 51 year-old male and reprogrammed via a non-integrating Sendai virus (KULSTEM iPSC core facility Leuven, Belgium, KSF-16-025). hiPSCs were cultured on Matrigel (Corning, #356237) in E8 flex medium (Thermo Fisher Scientific) supplemented with puromycin (0.5 µg/ml, Sigma Aldrich) and G418 (50 µg/ml, Sigma Aldrich) at 37°C/5% CO_2_. Medium was refreshed every 2-3 days and cells were passaged twice per week using an enzyme-free reagent (ReLeSR, Stem Cell Technologies).

HiPSCs were directly derived into Glutamatergic cortical layer 2/3 neurons and GABAergic neurons by overexpressing mouse neuronal determinant Neurogenin 2 (*Ngn2*) and Achaete-scute homolog 1 (*Ascl1*) upon doxycycline treatment^34, 67^, respectively as described in Pires Monteiro et al.^18^. Rodent astrocytes isolated from brain cortices of new-born (P1) Wistar rats were added in a 1:1 ratio at DIV 2. On DIV 3, the medium was replaced with Neurobasal medium (Thermo Fisher Scientific) supplemented with B-27 (Thermo Fisher Scientific), glutaMAX (Thermo Fisher Scientific), primocin (0.1 μg/ml, Invivogen), doxycycline (4 µg/ml, Sigma Aldrich), BDNF (10 ng/ml, Bioconnect), NT3 (10 ng/ml, Bioconnect) and forskolin (10 μM, Sigma Aldrich). Cytosine β-D-arabinofuranoside (Ara-C) (2 µM, Sigma Aldrich) was added to remove any proliferating cells. From this day onwards, half of the medium was refreshed 3 times a week. The medium was additionally supplemented with 2.5% FCS (Sigma Aldrich) from DIV 9 onwards. Doxycycline and forskolin were removed after DIV 13. Neuronal networks were kept at 37°C, 5% CO_2_ until the day of experiment.

### Experimental protocol

The experiments were conducted at 7 weeks *in vitro* in a computer-controlled incubator (temperature 37°C, 40% humidity, 5% CO_2_). After 30 minutes of accommodation in normoxia (oxygen 20%), cultures were exposed to 24 or 48 hours of hypoxia (oxygen 2%). Neuronal network activity was recorded with MEA for 10 minutes in normoxia and once every two hours in hypoxia. Cell viability was assessed in normoxia and after 24 and 48 hours of hypoxia.

During hypoxia, three experimental conditions reported below were used.

*1) Neuronal and astrocyte activation.* It was performed through optogenetic stimulation of excitatory neurons and astrocytes. To make excitatory neurons or astrocytes sensitive to blue light, 1 hour after plating (DIV 0 and DIV 2 for neurons and astrocytes, respectively) *Ngn2*-hiPSCs or astrocytes were transduced with an adeno-associated virus (AAV) serotype 2 encoding Channelrhodopsin-2 (ChR2) (AAV2-hSyn-hChR2(H134R)-mCherry, UNC Vector Core)^68^ and AAV serotype 2 encoding ChR2 (AAV-GFAP-hChR2(H134R)-mCherry, UNC Vector Core)^41, 69^, respectively. Afterwards, culture medium was replaced. On the day of the experiment, blue light (λ = 470 nm) was delivered to neuronal networks during the entire hypoxic period with the use of a Multiwell-Optogenetic prototype (Multi Channel Systems, Reutlingen, Germany) connected with the computer through the interface board. Light pulses (0.2 Hz, 10 mA for neuronal activation; 25 Hz, 0.3 mA for astrocyte activation) were applied for 3 minutes every 2 hours.
*2) Blockage of ANLS transporters.* Dihydrokainic acid (DHK) (Sigma-Aldrich) (i.e., EAAT2 selective inhibitor) was used at a final concentration of 300 μM^70^ and AR-C155858 (Sigma-Aldrich) (inhibitor of monocarboxylate transporters (MCTs) 1 and 2) was pre-diluted in DMSO and used at a final concentration of 1 μM^71^. The blockers were added in the medium independently before the onset of hypoxia.
*3) Addition of Lactate.* Sodium L-lactate and sodium D-lactate (Sigma Aldrich) were dissolved in Neurobasal medium (Thermo Fisher Scientific) and added before hypoxia at different concentrations (5-10-20 mM of L-lactate and 10 mM of D-lactate). Medium pH was evaluated by using a pH test paper Duotest pH 5.0-8.0 (Macherey-Nagel). We found that pH was not altered by lactate addition up to 20 mM (data not shown).

### MEAs recordings and analysis

Recordings of electrophysiological activity were performed with 24-well MEAs (Multichannel systems) (24 independent wells with 12 embedded microelectrodes, 30 μm in diameter and spaced 200 μm apart) covered with a Breathe Easier sealing membrane (Sigma Aldrich). Data were acquired with the Multiwell Screen software (Multi Channel Systems, Reutlingen, Germany) at a frequency of 10 kHz and signals were filtered between 100 and 3500 Hz. Data analysis was performed with the use of the Multiwell Analyzer software (Multi Channel Systems, Reutlingen, Germany) in combination with custom made MATLAB scripts (The Mathworks, Natick, MA, USA).

#### Spike detection

Spikes were detected if exceeding a threshold of 4.5 times the standard deviation of the baseline noise. An electrode was considered active if exhibiting at least 0.1 spikes/s. The mean firing rate (MFR) was calculated as the average of the number of spikes in time per electrode in a well, considering only the active electrodes.

#### Burst detection

Single channel bursts were detected when containing a minimum of 4 spikes with an inter spike interval between 50 ms and 100 ms. The minimum interval between bursts and burst duration were set at 100 ms and 50 ms, respectively. A channel was defined as bursting channel if exhibiting at least 0.4 burst/min.

#### Network burst detection

A network burst was defined as temporally overlapping single-channel bursts when the following qualifications apply: the number of distinct bursting channels needed to be at least 8 and at some time during the sequence of time at least 8 channels were simultaneously bursting (i.e., burst with temporal overlap > 0 ms with another burst). The network burst rate (NBR) was calculated as the number of network bursts detected in time in a well. The network burst duration (NBD) was calculated as the average duration of all network bursts detected in a well.

For each well, MFR, NBR and NBD were extracted for every timepoint and normalized with respect to the values from the normoxia phase. Afterwards, the average of normalized MFR, NBR and NBD was calculated for every timepoint among the wells with the same experimental condition. In experiments with neuronal and astrocyte activation, the parameters were evaluated only during the time windows in which stimulation was not delivered.

### Live/dead assays

CellEvent™ Caspase-3/7 Green Detection Reagent (Invitrogen) (1:500) was added to each well before the beginning of experiments, and incubated for 30 minutes at 37°C for detection of apoptotic cells (green). In neuronal networks not expressing ChR2, after the end of experiments Propidium Iodide (PI) (Invitrogen) (1:1000) was added to each well and incubated for 15 min at room temperature (RT) to detect dead cells (red). Then, coverslips were washed with PBS and fixated with 3.7% paraformaldehyde (PFA) for 15 min at RT. After washing with PBS, DAPI (Sigma Aldrich) (1:1000) was added into each well and incubated for 20 min at RT to stain cellular nuclei (blue). In neuronal networks expressing ChR2, apoptotic and dead cells were detected separately. Apoptotic cells were detected with CellEvent™ Caspase-3/7 Green Detection Reagent (Invitrogen) (1:500), as previously described. Dead cells were detected with a ReadyProbes™ Cell Viability Imaging Kit Blue/Green (Invitrogen): after the end of experiments, two drops per ml of NucBlue™ Live and NucGreen™ Dead reagents were added into each well and incubated for 15/30 min at RT to stain the nuclei of all cells (blue) and detect dead cells (green), respectively. Then, coverslips were washed with PBS and fixated with 3.7% paraformaldehyde (PFA) for 15 min at RT.

Coverslips were finally washed one more time with PBS and mounted with Mowiol on glass microscope slides. Coverslips were kept in the dark at RT for one night, then they were stored at 4°C. Epifluorescent pictures (n=8 per coverslip) were taken at a 40x magnification (0.085µm/pixel) with the use of a Nikon Eclipse 50i epifluorescence microscope (Nikon, Japan) and NIS-Elements Microscope Imaging Software (Nikon).

A custom made MATLAB script (The Mathworks, Natick, MA, USA) was used to set a fluorescence intensity threshold for each staining. Then, the total number of cells (DAPI or NucBlue™ Live positive, blue), the number of apoptotic cells (CellEvent™ Caspase-3/7 positive, green) and dead cells (PI positive, red, or both CellEvent™ Caspase-3/7 and PI positive, yellow, or NucGreen™ positive, green) were counted. The number of live cells was calculated by subtracting the apoptotic and dead cells to the total number of cells. The number of live, apoptotic and dead cells was considered as percentage of the total number of cells.

### Immunostaining

Neuronal networks were fixed with 3.7% paraformaldehyde (Sigma Aldrich) for 15 min at RT, washed with PBS (Sigma Aldrich) and stored at 4°C in PBS until stained. Samples were permeabilized with 0.2% Triton X-100 (Sigma Aldrich) in PBS for 5 min at RT, washed with PBS and blocked with 2% BSA (Sigma Aldrich) in PBS for 30 min at RT to block non-specific binding sites cells. The cultures were stained for rabbit anti-MAP2 (1:1000; Sigma M3696), rabbit anti-GFAP (1:500; Abcam ab7260) overnight at 4°C in blocking buffer. Samples were washed with PBS and stained with secondary antibodies for 1 h at RT, washed again and as a last step the nuclei were stained with DAPI (1:1000; Sigma Aldrich) 20 min at RT. Samples were washed and mounted with Mowiol (Sigma Aldrich). The secondary antibodies used were goat anti-mouse Alexa Fluor 488 (1:2000, Invitrogen A-11029) and goat anti-rabbit Alexa Fluor 568 (1:2000, Invitrogen A-11036). Images were taken at a 40× magnification with the use of a Nikon Eclipse 50i Epi-Fluorescence microscope (Nikon, Japan).

### Colorimetric assays

To measure lactate concentration in the culture medium, a Lactate Colorimetric Assay Kit (Abcam, ab65333) was used according to protocol of the manufacturer. Samples of medium were centrifuged, and supernatant was subsequently filtered over an 10 kD cut-off concentrator (Pierce, 88513) to remove proteins and debris. Results were analysed at specific light wavelengths in a Multiskan 60 (Thermo Scientific) plate reader. To make sure all read values were in the range of the calibration curve, series dilutions were made from the samples.

### Statistical analysis

Statistical analysis was performed with the use of GraphPad 9 (GraphPad Software, Inc., CA, USA). Normal distribution of data was ensured using the Kolmogorov-Smirnov normality test. Statistical analysis was performed with the parametric t-test or the non-parametric Mann Whitney test (for comparisons between two groups) and the two-way ANOVA test with post-hoc Bonferroni correction (for comparisons between more than two groups). To evaluate statistical significance p-values < 0.05 were considered to be significant. Exact p-values are reported in Supplementary Table S1-S4. Data are presented as mean ± standard error of the mean (SEM). The number of independent neuronal networks included for each experiment is reported in the figure legends. Graphs were created with the use of OriginLab 2019b (OriginLab Corporation).

## Acknowledgment

This research has been supported by an institutional research grant. Figures were prepared with BioRender.com

## Author contribution statement

M F conceived and supervised the study. M C, J M K, and M F designed all the experiments. M C and M L performed all experiments. M C performed data analysis. M F provided resources. M C, M J K and M F provides conceptualization and intellectual content. M C and M F wrote the paper. M L and J M K edited the paper.

## References

1. Donkor, E. S. Stroke in the 21st Century: A Snapshot of the Burden, Epidemiology, and Quality of Life. Stroke Res Treat 2018, (2018).

2. Hofmeijer, J. & Van Putten, M. J. A. M. Ischemic cerebral damage: An appraisal of synaptic failure. Stroke 43, 607–615 (2012).

3. Le Feber, J., Pavlidou, S. T., Erkamp, N., Van Putten, M. J. A. M. & Hofmeijer, J. Progression of neuronal damage in an in vitro model of the ischemic penumbra. PLoS One 11, 1–19 (2016).

4. Heiss, W. D. The ischemic penumbra: How does tissue injury evolve? Ann N Y Acad Sci 1268, 26–34 (2012).

5. Kleindorfer, D., Kissela, B. & Schneider, A. Eligibility for Recombinant Tissue Plasminogen Activator in Acute Ischemic Stroke. Stroke (2003).

6. Berkhemer, O. A. et al. A Randomized Trial of Intraarterial Treatment for Acute Ischemic Stroke. New England Journal of Medicine 372, 11–20 (2015).

7. Brouns, R. & De Deyn, P. P. The complexity of neurobiological processes in acute ischemic stroke. Clin Neurol Neurosurg 111, 483–495 (2009).

8. Auer, R. N. Neuroprotection in the Treatment of Brain Ischemia. Prog Cardiovasc Dis 271– 282 (2017).

9. Zhou, Z. et al. Advances in stroke pharmacology. Pharmacol Ther 191, 23–42 (2018).

10. Ginsberg, M. D. Neuroprotection for ischemic stroke: Past, present and future. Neuropharmacology 55, 363–389 (2008).

11. O’Collins, V. E. et al. 1,026 Experimental treatments in acute stroke. Ann Neurol 59, 467–477 (2006).

12. Moretti, A., Ferrari, F. & Villa, R. F. Neuroprotection for ischaemic stroke: Current status and challenges. Pharmacol Ther 146, 23–34 (2015).

13. Mao, Z., Bonni, A., Xia, F., Nadal-Vicens, M. & Greenberg, M. E. Neuronal activity-dependent cell survival mediated by transcription factor MEF2. Science (1979) 286, 785–790 (1999).

14. Gosh, A., Carnahan, J. & Greenberg, M. E. Requirement for BDNF in Activity-Dependent Survival of Cortical Neurons. Science (1979) 192, 263–265 (2016).

15. Ruijter, B. J. et al. Early electroencephalography for outcome prediction of postanoxic coma: A prospective cohort study. Ann Neurol 86, 203–214 (2019).

16. Taxis Di Bordonia E Valnigra, D., et al. The Association between Hypoxia-Induced Low Activity and Apoptosis Strongly Resembles That between TTX-Induced Silencing and Apoptosis. Int J Mol Sci 23, (2022).

17. Muzzi, L., Hassink, G. C., Levers, M., Hofmeijer, J. & Hospital, R. Mild stimulation improves neuronal survival in an in-vitro model of the ischemic penumbra. J Neural Eng (2019) doi:10.1088/1741-2552/ab51d4.

18. Pires Monteiro, S., et al. Neuroprotective effect of hypoxic preconditioning and neuronal activation in a in vitro human model of the ischemic penumbra. J Neural Eng 18, (2021).

19. Mosienko, V., Teschemacher, A. G. & Kasparov, S. Is L-lactate a novel signaling molecule in the brain? Journal of Cerebral Blood Flow and Metabolism 35, 1069–1075 (2015).

20. Magistretti, P. J. & Allaman, I. Lactate in the brain: From metabolic end-product to signalling molecule. Nat Rev Neurosci 19, 235–249 (2018).

21. Pellerin, L. & Magistretti, P. J. Sweet sixteen for ANLS. Journal of Cerebral Blood Flow and Metabolism 32, 1152–1166 (2012).

22. Pellerin, L. & Magistretti, P. J. Glutamate uptake into astrocytes stimulates aerobic glycolysis: A mechanism coupling neuronal activity to glucose utilization. Proc Natl Acad Sci U S A 91, 10625–10629 (1994).

23. Schurr, A., Payne, R. S., Miller, J. J. & Rigor, B. M. Brain lactate, not glucose, fuels the recovery of synaptic function from hypoxia upon reoxygenation: An in vitro study. Brain Res 744, 105–111 (1997).

24. Schurr, A., Payne, R. S., Miller, J. J. & Rigor, B. M. Brain lactate is an obligatory aerobic energy substrate for functional recovery after hypoxia: Further in vitro validation. J Neurochem 69, 423–426 (1997).

25. Cater, H. L., Chandratheva, A., Benham, C. D., Morrison, B. & Sundstrom, L. E. Lactate and glucose as energy substrates during, and after, oxygen deprivation in rat hippocampal acute and cultured slices. J Neurochem 87, 1381–1390 (2003).

26. Berthet, C. et al. Neuroprotective role of lactate after cerebral ischemia. Journal of Cerebral Blood Flow and Metabolism 29, 1780–1789 (2009).

27. Berthet, C., Castillo, X., Magistretti, P. J. & Hirt, L. New evidence of neuroprotection by lactate after transient focal cerebral ischaemia: Extended benefit after intracerebroventricular injection and efficacy of intravenous administration. Cerebrovascular Diseases 34, 329–335 (2012).

28. Schurr, A. & Gozal, E. Aerobic production and utilization of lactate satisfy increased energy demands upon neuronal activation in hippocampal slices and provide neuroprotection against oxidative stress. Front Pharmacol 3 JAN, 1–15 (2012).

29. Lam, T. I. et al. Intracellular pH reduction prevents excitotoxic and ischemic neuronal death by inhibiting NADPH oxidase. Proc Natl Acad Sci U S A 110, (2013).

30. Tauffenberger, A., Fiumelli, H., Almustafa, S. & Magistretti, P. J. Lactate and pyruvate promote oxidative stress resistance through hormetic ROS signaling. Cell Death Dis 10, (2019).

31. Yang, J. et al. Lactate promotes plasticity gene expression by potentiating NMDA signaling in neurons. Proc Natl Acad Sci U S A 111, 12228–12233 (2014).

32. Margineanu, M. B., Mahmood, H., Fiumelli, H. & Magistretti, P. J. L-Lactate Regulates the Expression of Synaptic Plasticity and Neuroprotection Genes in Cortical Neurons: A Transcriptome Analysis. Front Mol Neurosci 11, 1–17 (2018).

33. Banerjee, A., Ghatak, S. & Sikdar, S. K. l-Lactate mediates neuroprotection against ischaemia by increasing TREK1 channel expression in rat hippocampal astrocytes in vitro. J Neurochem 265–281 (2016) doi:10.1111/jnc.13638.

34. Mossink, B. et al. Cadherin-13 is a critical regulator of GABAergic modulation in human stem-cell-derived neuronal networks. Mol Psychiatry 27, 1–18 (2022).

35. Frega, M. et al. Neuronal network dysfunction in a model for Kleefstra syndrome mediated by enhanced NMDAR signaling. Nat Commun 10, 1–15 (2019).

36. Mossink, B. et al. Human neuronal networks on micro-electrode arrays are a highly robust tool to study disease-specific genotype-phenotype correlations in vitro. Stem Cell Reports 16, 2182– 2196 (2021).

37. Pellerin, L. & Magistretti, P. J. Glutamate uptake stimulates Na+, K+-ATPase activity in astrocytes via activation of a distinct subunit highly sensitive to ouabain. J Neurochem 69, 2132–2137 (1997).

38. Nedergaard, M. & Goldman, S. A. Carrier-mediated transport of lactic acid in cultured neurons and astrocytes. Am J Physiol Regul Integr Comp Physiol 265, (1993).

39. Ewaschuk, J. B., Naylor, J. M. & Zello, G. A. D-lactate in human and ruminant metabolism. Journal of Nutrition 135, 1619–1625 (2005).

40. Bozzo, L., Puyal, J. & Chatton, J. Y. Lactate Modulates the Activity of Primary Cortical Neurons through a Receptor-Mediated Pathway. PLoS One 8, 1–9 (2013).

41. Tang, F. et al. Lactate-mediated glia-neuronal signalling in the mammalian brain. Nat Commun 5, 1–13 (2014).

42. Schurr, A., Payne, R. S., Miller, J. J., Tseng, M. T. & Rigor, B. M. Blockade of lactate transport exacerbates delayed neuronal damage in a rat model of cerebral ischemia. Brain Res 895, 268–272 (2001).

43. Schurr, A., Miller, J. J., Payne, R. S. & Rigor, B. M. An increase in lactate output by brain tissue serves to meet the energy needs of glutamate-activated neurons. Journal of Neuroscience 19, 34–39 (1999).

44. Jourdain, P. et al. L-Lactate protects neurons against excitotoxicity: Implication of an ATP-mediated signaling cascade. Sci Rep 6, 1–13 (2016).

45. Ros, J., Pecinska, N., Alessandri, B., Landolt, H. & Fillenz, M. Lactate reduces glutamate-induced neurotoxicity in rat cortex. J Neurosci Res 66, 790–794 (2001).

46. Brunet, A., Datta, S. R. & Greenberg, M. E. Transcription-dependent and -independent control of neuronal survival by the PI3K-Akt signaling pathway. Curr Opin Neurobiol 11, 297–305 (2001).

47. Geng, X. et al. Synergetic neuroprotection of normobaric oxygenation and ethanol in ischemic stroke through improved oxidative mechanism. Stroke 44, 1418–1425 (2013).

48. Wang, Z. et al. Therapeutic potential of novel twin compounds containing tetramethylpyrazine and carnitine substructures in experimental ischemic stroke. Oxid Med Cell Longev 2017, (2017).

49. Zhang, G. et al. Neuroprotective Effect and Mechanism of Action of Tetramethylpyrazine Nitrone for Ischemic Stroke Therapy. Neuromolecular Med 20, 97–111 (2018).

50. Annoni, F. et al. Brain Protection after Anoxic Brain Injury: Is Lactate Supplementation Helpful ? Cells 1–11 (2021).

51. Vasile, F., Dossi, E. & Rouach, N. Human astrocytes: structure and functions in the healthy brain. Brain Struct Funct 222, 2017–2029 (2017).

52. Volterra, A. & Meldolesi, J. Astrocytes, from brain glue to communication elements: The revolution continues. Nat Rev Neurosci 6, 626–640 (2005).

53. Bélanger, M. & Magistretti, P. J. The role of astroglia in neuroprotection. Dialogues Clin Neurosci 11, 281–296 (2009).

54. Sofroniew, M. v. Molecular dissection of reactive astrogliosis and glial scar formation. Cell (2009) doi:10.1016/j.tins.2009.08.002.

55. Tuttolomondo, A., Raimondo, D. di, Sciacca, R., Pinto, A. & Licata, G. Inflammatory Cytokines in Acute Ischemic Stroke. Curr Pharm Des 3574–3589 (2008).

56. Endoh, M., Maiese, K. & Wagner, J. Expression of the inducible form of nitric oxide synthase by reactive astrocytes after transient global ischemia. Brain Res 651, 92–100 (1994).

57. Swanson, R. A., Ying, W. & Kauppinen, T. M. Astrocyte Influences on Ischemic Neuronal Death. Curr Mol Med 193–205 (2004).

58. Rosenberg, P. A. & Aizenman, E. Hundred-fold increase in neuronal vulnerability to glutamate toxicity in astrocyte-poor cultures of rat cerebral cortex. Neurosci Lett 103, (1989).

59. Chen, Y. et al. Astrocytes protect neurons from nitric oxide toxicity by a glutathione-dependent mechanism. J Neurochem 1601–1610 (2001).

60. Pablo, Y. de, Nilsson, M., Pekna, M. & Pekny, M. Intermediate filaments are important for astrocyte response to oxidative stress induced by oxygen – glucose deprivation and reperfusion. Histochem Cell Biol 81–91 (2013) doi:10.1007/s00418-013-1110-0.

61. Song, H., Stevens, C. F. & Gage, F. H. Astroglia induce neurogenesis from adult neural stem cells. Nature 39–44 (2002).

62. Mauch, D. H., Na, K. & Schumacher, S. CNS Synaptogenesis Promoted by Glia-Derived Cholesterol. Science (1979) 294, 1354–1357 (2001).

63. Christopherson, K. S. et al. Thrombospondins are astrocyte-secreted proteins that promote CNS synaptogenesis. Cell 120, 421–433 (2005).

64. Reperfusion, I. et al. Differential Regulation of Thrombospondin-1 and Thrombospondin-2 After Focal Cerebral. Stroke (2003) doi:10.1161/01.STR.0000047100.84604.BA.

65. Liu, Z. & Chopp, M. Astrocytes, therapeutic targets for neuroprotection and neurorestoration in ischemic stroke. Prog Neurobiol. vol. 176 139–148 (2016).

66. Choudhury, G. R. & Ding, S. Reactive astrocytes and therapeutic potential in focal ischemic stroke. Neurobiol Dis 234–244 (2017) doi:10.1016/j.nbd.2015.05.003.Reactive.

67. Frega, M. et al. Rapid neuronal differentiation of induced pluripotent stem cells for measuring network activity on micro-electrode arrays. Journal of Visualized Experiments 2017, 1–10 (2017).

68. Jin, L. et al. High-efficiency transduction and specific expression of ChR2opt for optogenetic manipulation of primary cortical neurons mediated by recombinant adeno-associated viruses. J Biotechnol 233, 171–180 (2016).

69. Gourine, A. V. et al. Astrocytes control breathing through pH-dependent release of ATP. Science (1979) 329, 571–575 (2010).

70. Arriza, J. L. et al. Functional Comparisons of Three Glutamate Cloned from Human Motor Cortex Transporter. J Neurosci 14, 5559–5569 (1994).

71. Ovens, M. J., Davies, A. J., Wilson, M. C., Murray, C. M. & Halestrap, A. P. AR-C155858 is a potent inhibitor of monocarboxylate transporters MCT1 and MCT2 that binds to an intracellular site involving transmembrane helices 7-10. Biochemical Journal 425, 523–530 (2010).

