## Supplementary information for "Neuroprotective role of lactate in a human *in vitro* model of the ischemic penumbra"

dr. Monica Frega

### Supplementary information

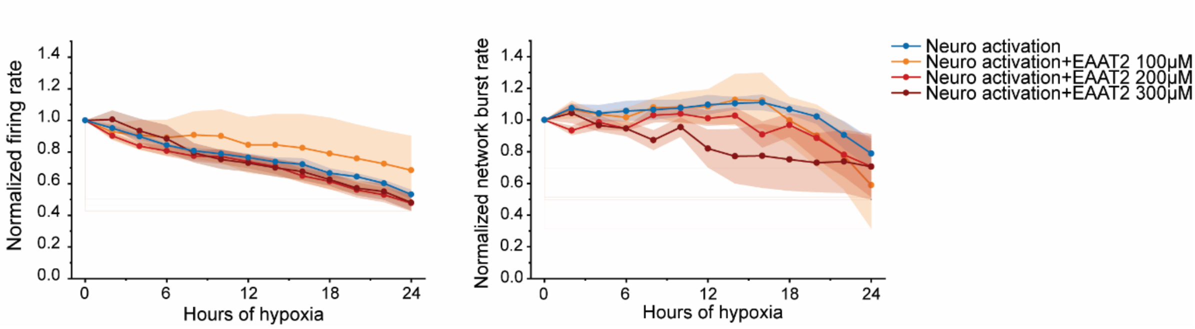


**Supplementary Figure 1.** Graphs showing the effect of EAAT2 blocker on MFR and NBR in neuronal networks activated with optogenetic stimulation. The values are normalized to the data of normoxia (neuro activation n=5, neuro activation + EAAT2 blocker 100 µM n=4, neuro activation + EAAT2 blocker 200 µM n=4, neuro activation + EAAT2 300 µM n=5). Two-way ANOVA test with multiple comparisons and post hoc Bonferroni correction was performed between conditions.

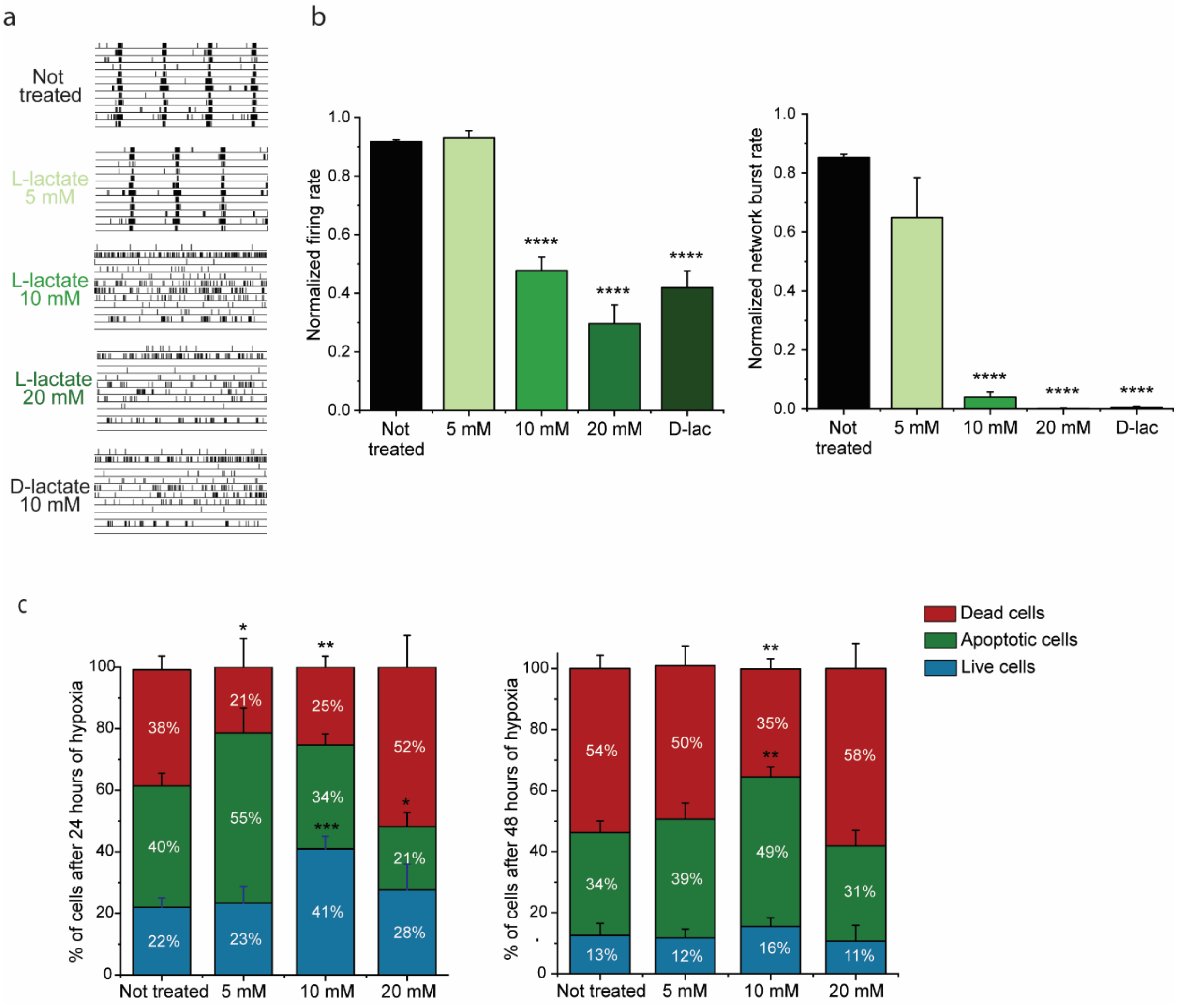


**Supplementary Figure 2. a** Representative raster plots showing 60 seconds of electrophysiological activity exhibited in normoxia by neuronal networks treated with different concentrations of lactate. **b** Bar graphs showing the effect of lactate addition on MFR and NBR in normoxia. The values are normalized to the data of the baseline phase before lactate addition (not treated n=7, L-lactate 2 mM n=5, L-lactate 5 mM n=5, L-lactate 10 mM n=8, L-lactate 20 mM n=5, D-lactate 10 mM n=2). ***$p<0.0001$, two-way ANOVA test and post hoc Bonferroni correction was performed between conditions. **c** Stacked column graphs showing the percentage of live , apoptotic and dead cells after 24 and 48 hours of hypoxia (not treated n=7, L-lactate 5 mM n=2, L-lactate 10 mM n=6, L-lactate 20 mM n=2). *$p<0.05$, **$p< 0.005$, ***$p<0.0005$, unpaired t test or Mann Whitney test was performed between conditions.

| Figure | Panel | Parameter and comparison | Hours of hypoxia | | p-value |
|---|---|---|---|---|---|
| ***1*** | ***e*** | ***Firing rate Not treated vs. neuro activation*** | 0 | ns | >0.9999 |
| | | | 2 | ns | 0.2642 |
| | | | 4 | ns | 0.4268 |
| | | | 6 | ns | 0.5173 |
| | | | 8 | ns | 0.6832 |
| | | | 10 | ns | 0.7915 |
| | | | 12 | ns | 0.5464 |
| | | | 14 | ns | 0.3790 |
| | | | 16 | ns | 0.2155 |
| | | | 18 | ns | 0.4034 |
| | | | 20 | ns | 0.3619 |
| | | | 22 | ns | 0.3882 |
| | | | 24 | ns | >0.9999 |
| | | ***Network burst rate Not treated vs. neuro activation*** | 0 | ns | >0.9999 |
| | | | 2 | **** | <0.0001 |
| | | | 4 | **** | <0.0001 |
| | | | 6 | **** | <0.0001 |
| | | | 8 | **** | <0.0001 |
| | | | 10 | **** | <0.0001 |
| | | | 12 | **** | <0.0001 |
| | | | 14 | **** | <0.0001 |
| | | | 16 | **** | <0.0001 |
| | | | 18 | **** | <0.0001 |
| | | | 20 | **** | <0.0001 |
| | | | 22 | **** | <0.0001 |
| | | | 24 | **** | <0.0001 |
| | ***g*** | ***Apoptotic cells Not treated vs. neuro activation*** | 24 | * | 0.0237 |
| | | ***Dead cells Not treated vs. neuro activation*** | 24 | ns | 0.0571 |
| | ***j*** | ***Firing rate Neuro activation vs. neuro activation + EAAT$_2$ blocker*** | 0 | ns | >0.9999 |
| | | | 2 | ns | >0.9999 |
| | | | 4 | ns | >0.9999 |
| | | | 6 | ns | >0.9999 |
| | | | 8 | ns | >0.9999 |
| | | | 10 | ns | >0.9999 |
| | | | 12 | ns | >0.9999 |
| | | | 14 | ns | >0.9999 |
| | | | 16 | ns | >0.9999 |
| | | | 18 | ns | >0.9999 |
| | | | 20 | ns | >0.9999 |
| | | | 22 | ns | >0.9999 |
| | | | 24 | ns | >0.9999 |
| | | ***Firing rate Neuro activation vs. neuro activation + MCT$_s$ blocker*** | 0 | ns | >0.9999 |
| | | | 2 | ns | >0.9999 |
| | | | 4 | ns | >0.9999 |
| | | | 6 | ns | >0.9999 |
| | | | 8 | ns | >0.9999 |
| | | | 10 | ns | >0.9999 |
| | | | 12 | ns | >0.9999 |
| | | | 14 | ns | >0.9999 |
| | | | 16 | ns | >0.9999 |
| | | | 18 | ns | >0.9999 |
| | | | 20 | ns | >0.9999 |
| | | | 22 | ns | >0.9999 |

| | | | | | |
|---|---|---|---|---|---|
| | | | 24 | ns | >0.9999 |
| | | ***Network burst rate Neuro activation vs. neuro activation + EAAT$_2$ blocker*** | 0 | ns | >0.9999 |
| | | | 2 | ns | >0.9999 |
| | | | 4 | ns | >0.9999 |
| | | | 6 | ns | >0.9999 |
| | | | 8 | ns | >0.9999 |
| | | | 10 | ns | >0.9999 |
| | | | 12 | ns | 0.9182 |
| | | | 14 | ns | 0.3882 |
| | | | 16 | ns | 0.3670 |
| | | | 18 | ns | 0.5107 |
| | | | 20 | ns | 0.7371 |
| | | | 22 | ns | >0.9999 |
| | | | 24 | ns | >0.9999 |
| | | ***Network burst rate Neuro activation vs. neuro activation + MCT$_s$ blocker*** | 0 | ns | >0.9999 |
| | | | 2 | ns | >0.9999 |
| | | | 4 | ns | >0.9999 |
| | | | 6 | ns | 0.3683 |
| | | | 8 | ns | >0.9999 |
| | | | 10 | ns | 0.5341 |
| | | | 12 | ns | 0.1161 |
| | | | 14 | * | 0.0444 |
| | | | 16 | ns | 0.0698 |
| | | | 18 | * | 0.0306 |
| | | | 20 | * | 0.0211 |
| | | | 22 | ** | 0.0088 |
| | | | 24 | ** | 0.0080 |

**Table S1**. Statistical analysis relative to Figure 1.

| Figure | Panel | Parameter and comparison | Hours of hypoxia | | p-value |
|---|---|---|---|---|---|
| ***2*** | ***c*** | ***Living cells Not treated vs. L-lactate*** | 24 | *** | 0.0002 |
| | | | 48 | ns | 0.0741 |
| | | ***Apoptotic cells Not treated vs. L-lactate*** | 24 | ns | 0.4464 |
| | | | 48 | ** | 0.0033 |
| | | ***Dead cells Not treated vs. L-lactate*** | 24 | ** | 0.0047 |
| | | | 48 | ** | 0.0010 |
| | | ***Living cells Not treated vs. D-lactate*** | 24 | ns | 0.1301 |
| | | | 48 | * | 0.0296 |
| | | ***Apoptotic cells Not treated vs. D-lactate*** | 24 | * | 0.0196 |
| | | | 48 | ns | 0.0532 |
| | | ***Dead cells Not treated vs. D-lactate*** | 24 | ns | 0.1047 |
| | | | 48 | ns | 0.8959 |
| | ***f*** | ***Lactate concentration Not treated vs. Astro activation*** | 0 | ** | 0.0017 |
| | ***h*** | ***Firing rate Not treated vs. astro activation*** | 0 | ns | >0.9999 |
| | | | 2 | **** | <0.0001 |
| | | | 4 | **** | <0.0001 |
| | | | 6 | **** | <0.0001 |
| | | | 8 | **** | <0.0001 |
| | | | 10 | **** | <0.0001 |
| | | | 12 | **** | <0.0001 |
| | | | 14 | **** | <0.0001 |
| | | | 16 | **** | <0.0001 |
| | | | 18 | **** | <0.0001 |
| | | | 20 | **** | <0.0001 |
| | | | 22 | **** | <0.0001 |
| | | | 24 | *** | 0.0002 |
| | ***i*** | ***Network burst rate Not treated vs. astro activation*** | 0 | ns | >0.9999 |
| | | | 2 | **** | <0.0001 |
| | | | 4 | **** | <0.0001 |
| | | | 6 | **** | <0.0001 |
| | | | 8 | **** | <0.0001 |
| | | | 10 | **** | <0.0001 |
| | | | 12 | **** | <0.0001 |
| | | | 14 | **** | <0.0001 |
| | | | 16 | **** | <0.0001 |
| | | | 18 | **** | <0.0001 |
| | | | 20 | **** | <0.0001 |
| | | | 22 | **** | <0.0001 |
| | | | 24 | **** | <0.0001 |
| | ***j*** | ***Network burst duration Not treated vs. astro activation*** | 0 | ns | >0.9999 |
| | | | 2 | ns | 0.1329 |
| | | | 4 | * | 0.0179 |
| | | | 6 | ** | 0.0028 |

| | | | | | |
|---|---|---|---|---|---|
| | | | 8 | ** | 0.0048 |
| | | | 10 | ** | 0.0027 |
| | | | 12 | *** | 0.0006 |
| | | | 14 | *** | 0.0001 |
| | | | 16 | *** | 0.0001 |
| | | | 18 | *** | 0.0007 |
| | | | 20 | *** | 0.0007 |
| | | | 22 | * | 0.0217 |
| | | | 24 | ns | 0.0766 |
| | ***g*** | ***Apoptotic cells Not treated vs. neuro activation*** | 24 | ** | 0.0031 |
| | | ***Dead cells Not treated vs. neuro activation*** | 24 | ns | 0.8383 |

**Table S2**. Statistical analysis relative to Figure 2.

| Figure | Panel | Parameter and comparison | Hours of hypoxia | | p-value |
|---|---|---|---|---|---|
| ***S1*** | | ***Firing rate*** ***Neuro activation vs. neuro activation + EAAT$_2$ blocker 100µM*** | 0 | ns | >0.9999 |
| | | | 2 | ns | >0.9999 |
| | | | 4 | ns | >0.9999 |
| | | | 6 | ns | >0.9999 |
| | | | 8 | ns | >0.9999 |
| | | | 10 | ns | >0.9999 |
| | | | 12 | ns | >0.9999 |
| | | | 14 | ns | >0.9999 |
| | | | 16 | ns | >0.9999 |
| | | | 18 | ns | >0.9999 |
| | | | 20 | ns | >0.9999 |
| | | | 22 | ns | >0.9999 |
| | | | 24 | ns | >0.9999 |
| | | ***Firing rate*** ***Neuro activation vs. neuro activation + EAAT$_2$ blocker 200µM*** | 0 | ns | >0.9999 |
| | | | 2 | ns | >0.9999 |
| | | | 4 | ns | >0.9999 |
| | | | 6 | ns | >0.9999 |
| | | | 8 | ns | >0.9999 |
| | | | 10 | ns | >0.9999 |
| | | | 12 | ns | >0.9999 |
| | | | 14 | ns | >0.9999 |
| | | | 16 | ns | >0.9999 |
| | | | 18 | ns | >0.9999 |
| | | | 20 | ns | >0.9999 |
| | | | 22 | ns | >0.9999 |
| | | | 24 | ns | >0.9999 |
| | | ***Firing rate*** ***Neuro activation vs. neuro activation + EAAT$_2$ blocker 300µM*** | 0 | ns | >0.9999 |
| | | | 2 | ns | >0.9999 |
| | | | 4 | ns | >0.9999 |
| | | | 6 | ns | >0.9999 |
| | | | 8 | ns | >0.9999 |
| | | | 10 | ns | >0.9999 |
| | | | 12 | ns | >0.9999 |
| | | | 14 | ns | >0.9999 |
| | | | 16 | ns | >0.9999 |
| | | | 18 | ns | >0.9999 |
| | | | 20 | ns | >0.9999 |
| | | | 22 | ns | >0.9999 |
| | | | 24 | ns | >0.9999 |
| | | ***Network burst rate*** ***Neuro activation vs. neuro activation + EAAT$_2$ blocker 100µM*** | 0 | ns | >0.9999 |
| | | | 2 | ns | >0.9999 |
| | | | 4 | ns | >0.9999 |
| | | | 6 | ns | >0.9999 |
| | | | 8 | ns | >0.9999 |
| | | | 10 | ns | >0.9999 |
| | | | 12 | ns | >0.9999 |
| | | | 14 | ns | >0.9999 |
| | | | 16 | ns | >0.9999 |
| | | | 18 | ns | >0.9999 |
| | | | 20 | ns | >0.9999 |
| | | | 22 | ns | >0.9999 |
| | | | 24 | ns | >0.9999 |
| | | ***Network burst rate*** ***Neuro activation vs. neuro activation + EAAT$_2$ blocker 200µM*** | 0 | ns | >0.9999 |
| | | | 2 | ns | >0.9999 |
| | | | 4 | ns | >0.9999 |
| | | | 6 | ns | >0.9999 |

| | | | | | |
|---|---|---|---|---|---|
| | | | 8 | ns | >0.9999 |
| | | | 10 | ns | >0.9999 |
| | | | 12 | ns | >0.9999 |
| | | | 14 | ns | >0.9999 |
| | | | 16 | ns | 0.6079 |
| | | | 18 | ns | >0.9999 |
| | | | 20 | ns | >0.9999 |
| | | | 22 | ns | >0.9999 |
| | | | 24 | ns | >0.9999 |
| | | ***Network burst rate Neuro activation vs. neuro activation + EAAT$_2$ blocker 300µM*** | 0 | ns | >0.9999 |
| | | | 2 | ns | >0.9999 |
| | | | 4 | ns | >0.9999 |
| | | | 6 | ns | >0.9999 |
| | | | 8 | ns | >0.9999 |
| | | | 10 | ns | >0.9999 |
| | | | 12 | ns | 0.9182 |
| | | | 14 | ns | 0.3882 |
| | | | 16 | ns | 0.3670 |
| | | | 18 | ns | 0.5107 |
| | | | 20 | ns | 0.7371 |
| | | | 22 | ns | >0.9999 |
| | | | 24 | ns | >0.9999 |

**Table S3**. Statistical analysis relative to Supplementary Figure 1.

| Figure | Panel | Parameter and comparison | Hours of hypoxia | | p-value |
|---|---|---|---|---|---|
| ***S2*** | ***b*** | ***Firing rate*** ***Not treated vs. 5 mM*** | 0 | ns | >0.9999 |
| | | ***Firing rate*** ***Not treated vs. 10 mM*** | 0 | **** | <0.0001 |
| | | ***Firing rate*** ***Not treated vs. 20 mM*** | 0 | **** | <0.0001 |
| | | ***Firing rate*** ***Not treated vs. D-lac*** | 0 | **** | <0.0001 |
| | | ***Network burst rate*** ***Not treated vs. 5 mM*** | 0 | ns | 0.2919 |
| | | ***Network burst rate*** ***Not treated vs. 10 mM*** | 0 | **** | <0.0001 |
| | | ***Network burst rate*** ***Not treated vs. 20 mM*** | 0 | **** | <0.0001 |
| | | ***Network burst rate*** ***Not treated vs. D-lac*** | 0 | **** | <0.0001 |
| | ***c*** | ***Living cells*** ***Not treated vs. 5 mM*** | 24 | ns | 0.7874 |
| | | | 48 | ns | 0.0817 |
| | | ***Apoptotic cells*** ***Not treated vs. 5 mM*** | 24 | ns | 0.0516 |
| | | | 48 | ns | 0.5184 |
| | | ***Dead cells*** ***Not treated vs. 5 mM*** | 24 | * | 0.0188 |
| | | | 48 | ns | 0.7090 |
| | | ***Living cells*** ***Not treated vs. 10 mM*** | 24 | *** | 0.0002 |
| | | | 48 | ns | 0.0741 |
| | | ***Apoptotic cells*** ***Not treated vs. 10 mM*** | 24 | ns | 0.4464 |
| | | | 48 | ** | 0.0033 |
| | | ***Dead cells*** ***Not treated vs. 10 mM*** | 24 | ** | 0.0047 |
| | | | 48 | ** | 0.0010 |
| | | ***Living cells*** ***Not treated vs. 20 mM*** | 24 | ns | 0.7303 |
| | | | 48 | ns | 0.2700 |
| | | ***Apoptotic cells*** ***Not treated vs. 20 mM*** | 24 | * | 0.0497 |
| | | | 48 | ns | 0.7557 |
| | | ***Dead cells*** ***Not treated vs. 20 mM*** | 24 | ns | 0.1885 |
| | | | 48 | ns | 0.6470 |

**Table S4**. Statistical analysis relative to Supplementary Figure 2.
